# Stage- and tissue-specific gene editing using 4-OHT inducible Cas9 in whole organism

**DOI:** 10.1101/2025.07.07.663418

**Authors:** Yaqi Li, Weiying Zhang, Zihang Wei, Han Li, Xin Liu, Tao Zheng, Tursunjan Aziz, Cencan Xing, Anming Meng, Xiaotong Wu

**Affiliations:** Laboratory of Molecular Developmental Biology, State Key Laboratory of Membrane Biology, Tsinghua-Peking Center for Life Sciences, School of Life Sciences, Tsinghua University, Beijing 100084, China; Guangzhou Laboratory, Guangzhou 510320, Guangdong Province, China

**Keywords:** CRISPR/Cas9, Estrogen receptor, Conditional knockout, Germ cells, Zebrafish

## Abstract

Vertebrate genes usually function in specific tissues at specific stages, and thus their functional studies require conditional knockout or editing. In the zebrafish, spatiotemporally inducible genome editing, especially during early embryonic development, is still challenging. Here, we report inducible Cas9-based gene editing in specific cell types at desired stages in zebrafish. We show that the nCas9^ERT2^ fusion protein consisting of Cas9 and estrogen receptor flanked by two nuclear localization signals is normally located in the cytoplasm and efficiently translocated into the nucleus upon 4-hydroxytamoxifen (4-OHT) treatment in cultured cells or embryos. As a proof-of-concept, we demonstrate that target genes in primordial germ cells or germ cells of transgenic zebrafish embryos or adults with stable expression of nCas9^ERT2^ and gene-specific guide RNAs (gRNAs) can be induced to mutate by application of 4-OHT. This inducible nCas9^ERT2^ system also works in mouse early embryos. Thus, this inducible gene editing approach will be another choice for studying the temporospatial function of genes at the organismal level. Hence, the inducible nCas9^ERT2^ system expands the tissue- and stage-specific gene editing toolkit.

## Introduction

The boom in genome editing technologies in recent years, specifically those using the CRISPR (Clustered Regularly Interspaced Short Palindromic Repeats)/CRISPR-associated (Cas) system, has helped simplify the process of generating genetically engineered animal models (1–6). However, traditional whole-body knock-out in early embryos via zygotic injection of Cas9 and gRNAs makes it inapplicable to elucidate cell type- and developmental stage-specific functions of genes. Thus, inducible conditional knock-out technologies are greatly desired in these circumstances. Much efforts have been made previously to improve traditional CRISPR/Cas9 technology to achieve cell- or tissue-specific genome editing at the organismal level by driving Cas9 expression under a tissue-specific promoter (7–12) or using a system that works only in particular biological contexts. For example, combined with Gal4/UAS binary system (13–17) or Cre/loxP system (18–20), Cas9 expression could be controlled by tissue-specific Gal4 transcription factor or Cre recombinase, in which UAS (upstream activation sequence) and the Stop cassette have been placed upstream of the Cas9 coding sequence, respectively. On the other hand, temporally controlled genome editing technologies have been applied more to cultured cells. For example, Cas9 expression can be repressed by TetR (tet repressor) protein via TetO (tetracycline operator) and activated when DOX (doxycycline) treatment via blocking TetR (21). In addition, assembling of the split Cas9 protein fragments induced by small molecules or light (22, 23) or alteration of the Cas9 protein conformation triggered by small molecules (24, 25) have also been applied to achieve temporal control of genome editing. However, due to uncommon specialized equipment, the complexity of the expression systems, or limited cell-permeability of inducible drugs, only a few of these inducible CRISPR/Cas9 systems have been utilized *in vivo* (21, 26).

In mice, inducible tissue-specific genome editing technologies based on Cre recombinase were widely used, such as TetO-Cre induced by Dox (27–30) and Cre variant with estrogen receptor (Cre-ER) induced by xenoestrogens tamoxifen (31–34). The subcellular localization and activity of the Cre recombinase can be regulated by tamoxifen or a related analog, such as 4-OHT (4-hydroxytamoxifen) when Cre is linked with a mutated ligand-binding domain of estrogen receptor (ERT2) to avoiding endogenous estrogen interference (32, 35). Notably, the Cre-based system requires the knock-in of two loxP fragments into the target gene locus (36–38). Therefore, the popularity of the inducible Cre/loxP system in zebrafish is largely impeded by low knock-in germline transmission rate in zebrafish (39–42). Moreover, fusion of ERT2 to Cas9 triggers nuclear translocation of Cas9 protein upon 4-OHT treatment, which has been demonstrated in mammalian cell lines (43, 44). Thus, we speculated that Cas9-ERT2 might also render highly efficient and convenient spatiotemporal gene editing in zebrafish.

Early development of zebrafish embryos is regulated by many maternally expressed genes (45, 46), some of which are also expressed zygotically. It has been observed that zygotic mutants of some maternal/zygotic genes are embryonic lethal, making it difficult to obtain their maternal mutants for investigating functions of their maternal products (47). Regarding germ cells (GCs), their development involves interaction with surrounding somatic cells, in which a single gene may express in both cell types thus elucidation of its function in a specific cell type is required. In this study, therefore, we made effort to establish an inducible gene knockout system, based on CRISPR/Cas9 technology, to achieve temporal gene knockout in zebrafish GCs.

## Results

### Design and test of 4-OHT inducible nucleus-localizing Cas9

To establish a 4-OHT inducible Cas9 system, we made several pCS2 vector-based expression constructs using zebrafish codon-optimized Cas9 (48), ERT2, SV40 nuclear localization signal (NLS1) and nucleoplasmin NLS (NLS2) (Fig. S1A). After transfection into HEK293T cells, we found that only Cas9-NLS1-ERT2-NLS2 protein (nCas9^ERT2^) showed nuclear transport upon 5 μM 4-OHT treatment (Fig. 1A, and S1B), suggesting the importance of NLS positioning in the fusion protein for its inducible nuclear translocation. To test the ability of nCas9^ERT2^ to edit target genes, its expression vector and the plasmid *pU6a-EMX1-gRNA* for targeting human *EMX1* locus (2) were cotransfected into HEK293T cells. T7 endonuclease I (T7EI) assay revealed that the *EMX1* locus was successfully edited only after 4-OHT treatment, which was similar to the editing effect by expressing nCas9 and *EMX1* gRNA from a single plasmid (Fig. 1B). Therefore, nCas9^ERT2^ was used in subsequent studies.

**Fig. 1.**
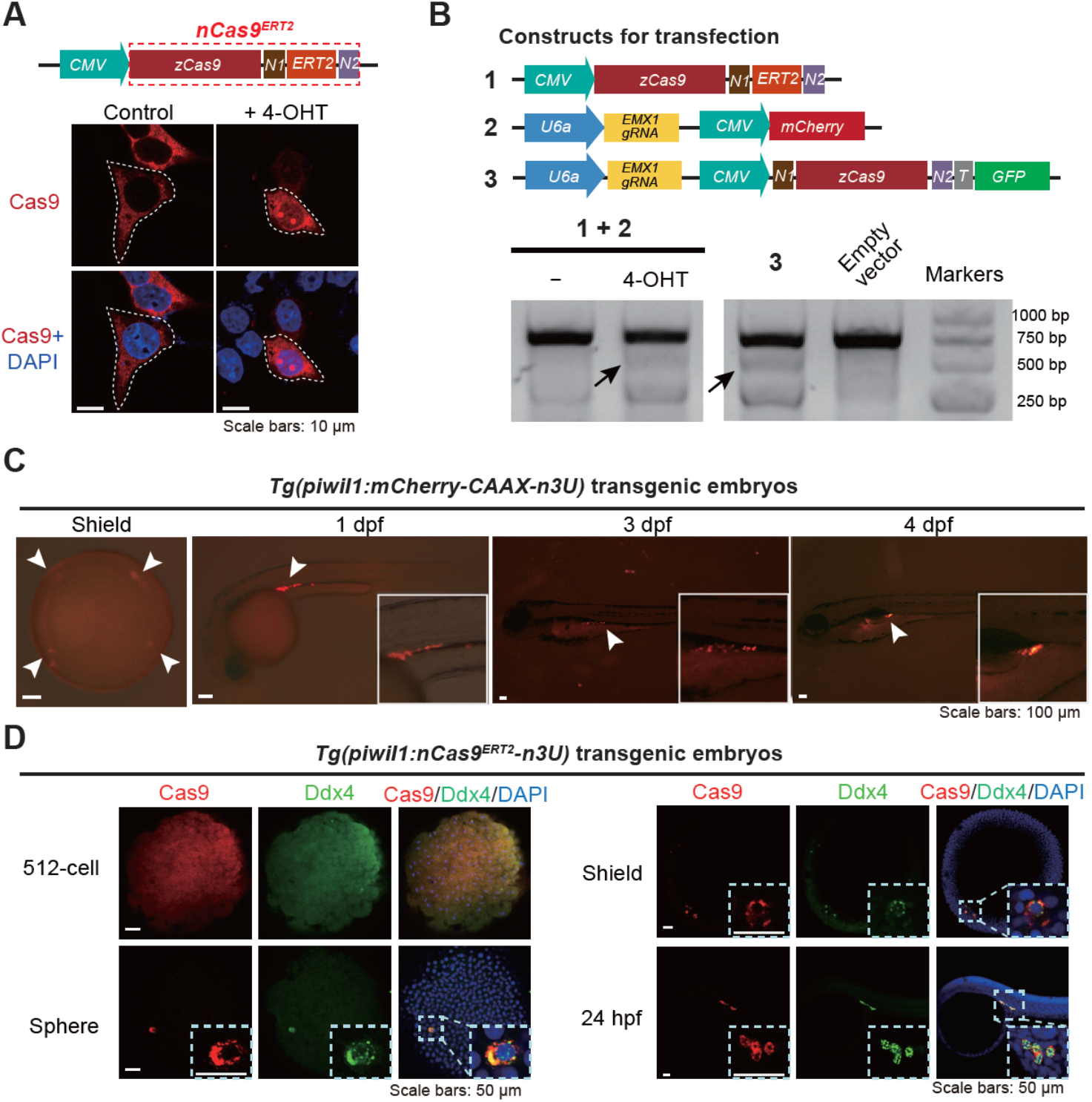
Nuclear translocation and gene editing effectiveness of nCas9^ERT2^. (A) 4-OHT induced nuclear translocation of nCas9^ERT2^ in human HEK293T cells. Cells were transfected by an expression construct (top) and immunostained with Cas9 antibody and DAPI staining (bottom). The shown were representative images with Cas9-positive cells outlined by the dashed line. N1, SV40 NLS; N2, nucleoplasmin NLS. (B) Gene editing effectiveness of 4-OHT induced nCas9^ERT2^ in HEK293T cells. Endogenous *EMX1* was the target. Top, compositions of constructs with designated numbers for transfection (T, T2A); bottom, gel electrophoretic image showing T7E1 assay outcome of *EMX1* locus. The mutant band was indicated by arrows. (C) The indicated transgenic embryos express mCherry specifically in PGCs (indicated by arrowheads), confirming the PGC specificity of the zebrafish *piwil1* promoter. Inserts showed arrowhead-pointed areas. (D) Expression pattern of nCas9^ERT2^ in indicated transgenic embryos at different stages. Embryos were immunostained with Cas9 and Ddx4 (a PGC-specific marker) antibodies plus DAPI staining. Note that Cas9 started to specifically express in PGCs from the sphere stage onwards. Inserts showed enlarged areas harboring PGCs.

### 4-OHT induced nuclear translocation of nCas9^ERT2^ in zebrafish PGCs

The zebrafish *piwil1* (also called *ziwi*) has been reported to be expressed in primordial germ cells (PGCs) in early embryos and in oogonia and oocytes in adults (49, 50). Based on previously reported 4.8-kb (kilo-base pairs) *piwil1* promoter with germline transcriptional specificity (51), promoter dissection by transient transgenesis in zebrafish identified its 2.5-kb proximal region with transgenic activity in PGCs/GCs (Fig. S2A). This 2.5-kb *piwil1* promoter was used subsequently. To enhance the germ cell expression specificity, as the 3’ untranslated region (3’UTR) sequence of *nanos3* (previously named *nanos1*) mRNA is crucial for PGC location (52–55)), we made the *Tol2* transposon-based construct *Tol2 (piwil1:mCherry-CAAX-n3U)* by combining the 2.5-kb *piwil1* promoter with the *nanos3* 3’UTR (n3U) to enhance GC-specific localization of expressed target mRNA. In resulted *Tg(piwil1:mCherry-CAAX-n3U)* germline transgenic embryos, the mCherry fluorescence, as observed by fluorescent microscopy, was clearly restricted to PGCs/GCs from 6 hours postfertilization (hpf) (shield stage) to 4 days postfertilization (dpf) (Fig. 1C). However, immunofluorescence using these embryos detected mCherry signal in all cells at the 512-cell stage (2.75 hpf), mainly in PGCs at the sphere stage (4 hpf), and then exclusively in PGCs thereafter (Fig. S2B).

We utilized the 2.5-kb *piwil1* promoter, *nCas9*^*ERT2*^, and n3U to generate the zebrafish *Tg(piwil1:nCas9n*^*ERT2*^*-n3U)* transgenic line. Immunofluorescence observation indicated that, in *Tg(piwil1:nCas9n*^*ERT2*^*-n3U)* transgenic embryos, Cas9 protein was ubiquitously distributed, and became localized in Ddx4/Vasa-positive PGCs at and after the sphere stage (Fig. 1D). Importantly, nCas9^ERT2^ protein was predominantly present in the cytoplasm. Then, we explored the way to induce nuclear transport of nCas9^ERT2^ in *Tg(piwil1:nCas9n*^*ERT2*^*-n3U)* embryos. These embryos were incubated in 20 μM 4-OHT at normal raising buffer and temperature kept out of the light from 4 hpf onward and collected at different time points in an interval of 2 hours (Fig. 2A) for Cas9 immunofluorescence. Results indicated that nCas9^ERT2^ protein remained in the cytoplasm of PGCs at 1 hour post-treatment (hpt), became detectable in the nucleus at 2 hpt, and was highly enriched in the nucleus at 4 and 6 hpt (Fig. 2B). Thus, nuclear transport of nCas9^ERT2^ can be achieved in PGCs of living embryos by a relatively shorter period of 4-OHT treatment. Besides, we did not see an adverse effect of 4-OHT on embryonic development in our treating conditions.

**Fig. 2.**
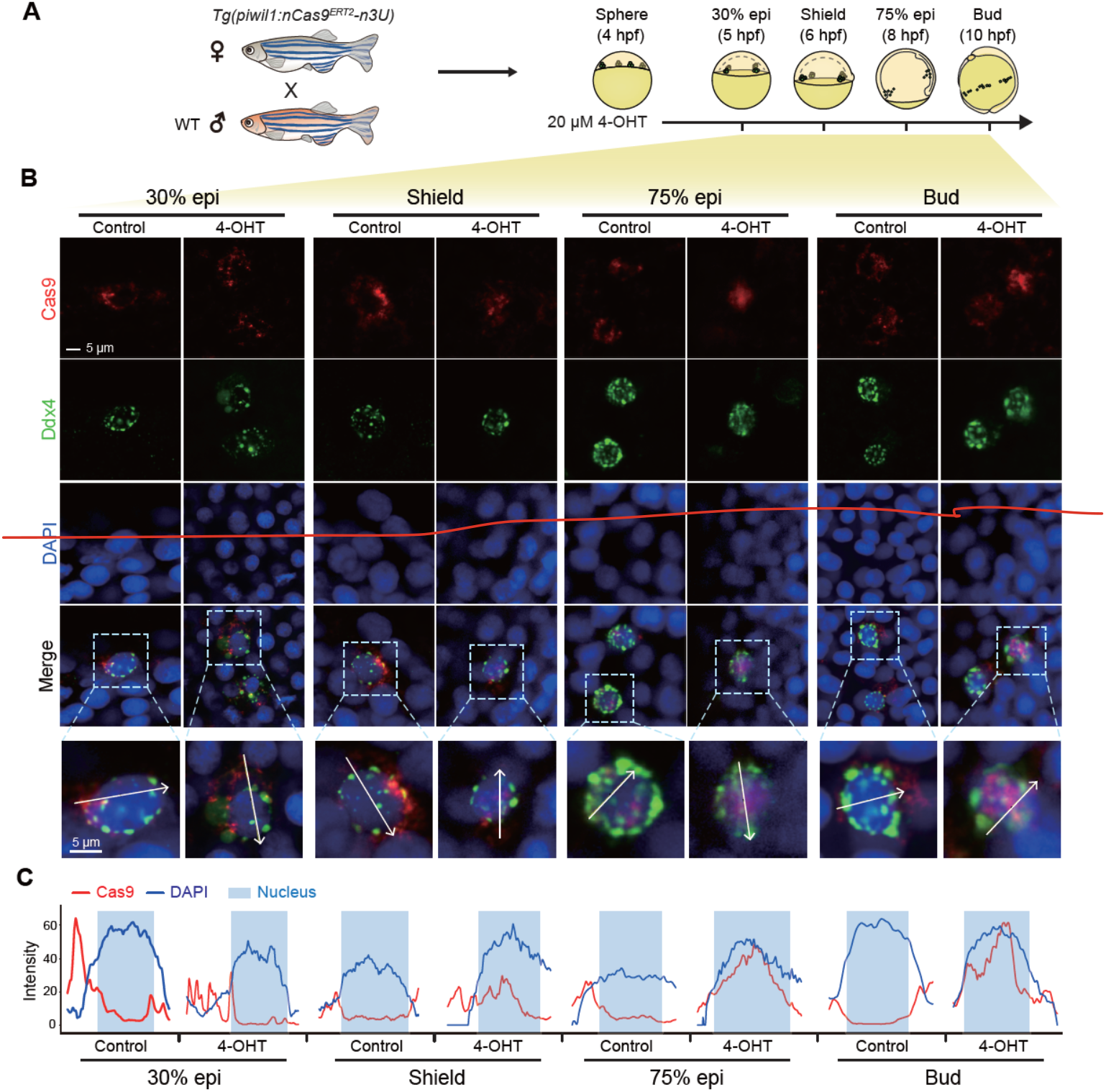
4-OHT induced nuclear translocation of PGC-specific nCas9^ERT2^ in transgenic zebrafish embryos. (A) Schematics showing the mating strategy and 4-OHT treatment of embryos. (B) Immunostaining results for Cas9 (red) and Ddx4 (green) with nuclei stained by DAPI before and after 4-OHT treatment. The dashed boxes in the fourth row were enlarged and shown in the fifth row. (C) Curve plots showing the fluorescence intensity measurements of Cas9 (red) and DAPI (blue) along the white arrow in the fifth row in (C) The blue boxes indicated regions of the nucleus.

### Stage-specific editing of *tbxta* gene in PGCs of embryos by the inducible nCas9^ERT2^ system

To test whether 4-OHT is able to induce sufficient amount of nuclear nCas9^ERT2^ to execute stage-specific gene editing in *Tg(piwil1:nCas9n*^*ERT2*^*-n3U)* transgenic embryos, we chose to target the *tbxta*/*ntl* gene, which is zygotically expressed in mesendodermal progenitors and the notochord (56). We adopted two strategies: 1) injection of synthetic *tbxta*-gRNA and *mCherry-n3U* mRNA (for labeling PGCs) into one-cell stage *Tg(piwil1:nCas9-n3U)* transgenic embryos, in which nCas9 was not fused to ERT2 and thus could transport to the nucleus without 4-OHT induction (Fig. 3A, left); 2) *Tg(piwil1:nCas9*^*ERT2*^*-n3U)* transgenic embryos injected with *tbxta*-gRNA and *mCherry-n3U* mRNA at the one-cell stage and treated with 40 μM 4-OHT from the sphere stage onwards (Fig. 3A, right). The resulted embryos were observed at 24 hpf, followed by identification of mutated alleles of *tbxta* using isolated mCherry-positive PGCs and mCherry-negative somatic cells (Fig. 3A). In case of *Tg(piwil1:nCas9-n3U)* embryos injected by *tbxta*-gRNA, 28.9% (30/104) exhibited truncation of posterior trunk and lack of the notochord at 24 hpf (class III), which phenocopied *tbxta* mutants (57, 58), 64.4% (67/104) had mildly truncated posterior trunk (class II), and the remaining had a normal trunk (class I) (Fig. 3B). Cloning and sequencing of the *tbxta-gRNA* targeted region indicated that 70.6% (12/17) and 50.0% (10/20) of alleles were mutated in mCherry-positive PGCs and mCherry-negative somatic cells in class III embryos at 24 hpf (Fig. 3C and D), respectively. In case of *tbxta*-gRNA injected and 4-OHT treated *Tg(piwil1:nCas9*^*ERT2*^*-n3U)* embryos, all (n=30) showed normal morphology at 24 hpf; however, 34.6% of *tbxta* alleles were mutated in PGCs whereas no mutated alleles were found in somatic cells (Fig. 3C and D). These results indicated that gene editing activity of nCas9^ERT2^ can be temporally controlled during embryogenesis through the application of 4-OHT. Importantly, stage- and PGCs-specific gene editing by using our system could allow loss-of-function investigation of a maternal gene if its zygotic mutations cause lethality.

**Fig. 3.**
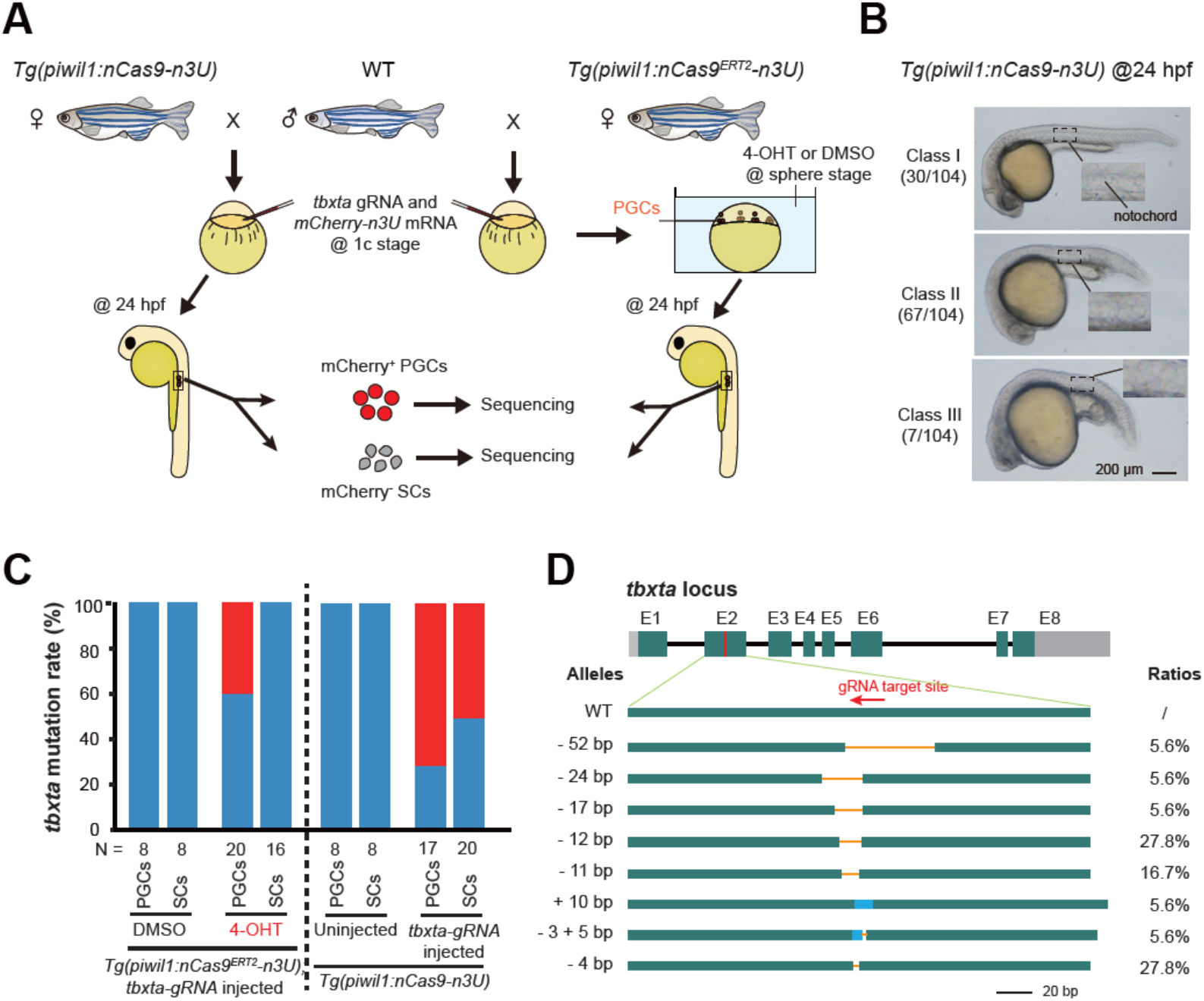
Stage- and PGC-specific mutation of *tbxta* locus in zebrafish nCas9 transgenic embryos. (A) Schematics of experimental procedures. *Tg(piwil1:nCas9-n3U)* (left) or *Tg(piwil1:nCas9*^*<sup>ERT2</sup>*^*-n3U)* (right) transgenic females were crossed to WT males, which produced embryos with maternal expression of non-inducible nCas9 or inducible nCas9^ERT2^, respectively. SCs, somatic cells. (B) Morphology of 24-hpf stage *Tg(piwil1:nCas9-n3U)* embryos injected with synthetic *tbxta-gRNA* at one-cell stage. The ratio of embryos with shown representative morphology was indicated in parentheses. The boxed areas were enlarged in the inserts. (C) Mutation rate of *tbxta* locus in mCherry-positive PGCs or mCherry negative SCs in different groups at 24 hpf. A region in the *tbxta* locus was amplified and cloned, followed by sequencing individual clones. N, number of sequenced clones. (D) Mutant types and their ratios at the *tbxta* locus. E1-E8, exons; green, WT sequence; gray, untranslated regions; orange line, deleted region; blue, inserted region.

### GC-specific editing of *huluwa* gene in young fish by the inducible nCas9^ERT2^ system

To test the utility of the inducible nCas9^ERT2^ system in adults, we chose to target the *huluwa* (*hwa*) gene, which is a maternal gene and its maternal mutants lack the dorsal organizer and body axis (59). To this end, we generated the *Tg(U6:hwa-gRNAs;ea:GFP)* transgenic line, which simultaneously expresses two previously reported *huluwa* gRNAs (59) as well as GFP for easy identification of transgenic fish (Fig. 4A). By crossing *Tg(piwil1:nCas9*^*ERT2*^*-n3U)* fish with *Tg(U6:hwa gRNAs;ef1α:GFP)* fish, we obtained *Tg(piwil1:nCas9*^*ERT2*^*-n3U;U6:hwa-gRNAs;ef1a:GFP)* double transgenic zebrafish. When grown to 45 dpf, these fish were treated twice, each lasting 12 hours with an interval of 12 hours for resting, with 5 μM 4-OHT, followed by raising in normal conditions and keeping out of the light. After reaching sexual maturity (about 3 months post fertilization (mpf)), the treated females were picked up to mate with wild-type (WT) males to produce embryos (Fig. 4B). Then the effectiveness of mutations in GCs could be evaluated by dorsal-ventral axis formation defects as reported previously in maternal *hwa* mutants (59). We observed that 1.2 - 8.6% of embryos derived from the transgenic females treated with 4-OHT could give birth to severely ventralized embryos mimicking M*hwa* mutants (Fig. 4C), which implies that both alleles of *hwa* in some germ stem cells have been mutated to lose function. It is worth noting that the ratio of ventralized embryos was low and varied among individuals. Further optimization of 4-OHT treatments may be needed to achieve a higher gene editing efficiency. By sequencing *hwa* alleles in individual ventralized embryos, we detected different types of mutations, among which the one-base insertion was the predominant type (Fig. 4D). Nevertheless, these results support the idea that our inducible nCas9^ERT2^ system works in adult fish.

**Fig. 4.**
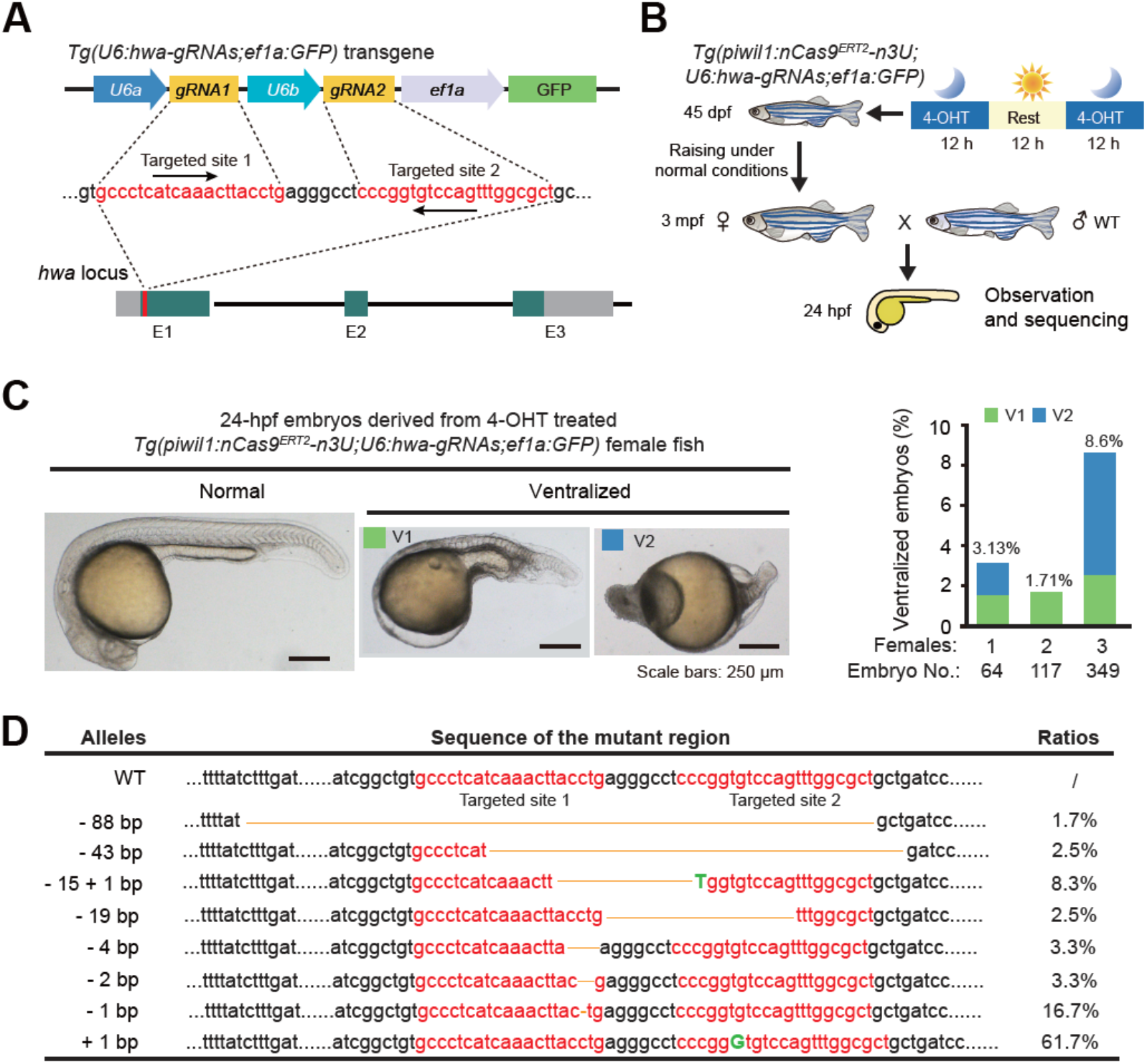
4-OHT induced *hwa* mutations in GCs of young females. (A) Zebrafish *hwa* locus and *hwa-gRNAs* transgene structures. E1-E3, exons; gray bars, untranslated region. (B) Procedure for 4-OHT treatments of the double transgenic young fish and subsequent mating and embryonic analysis. Moon, night; Sun, daytime. (C, D) Morphology of representative embryos at 24 hpf (left) and ratio of ventralized embryo types (right bar graph). (D) Mutant alleles at the *hwa* locus in ventralized embryos. A region spanning two target sides was amplified and cloned for sequencing. Dotted line, bases not shown; orange line, deleted bases; red letters, bases in gRNAs; green capital letters, inserted bases.

### Inducible nCas9^ERT2^ system helps determine implication of PGCs-expressed *tbx16* in PGC migration

The zebrafish *tbx16* is initially found to affect movements of mesodermal precursors and somite formation (60, 61). It is observed later on that *tbx16* mutants also exhibit abnormal locations of PGCs outside the gonad regions (62), and *tbx16* is also expressed in PGC as early as the dome stage (4.3 hpf) (63). This raises a puzzling question: defective PGC migration in *tbx16* mutant is ascribed to loss of Tbx16 in somatic cells or in PGCs. We thought that our nCas9^ERT2^ system might help address this question.

Then, we generated the *Tg(U6:tbx16-gRNAs;cryaa:CFP)* transgenic line that express two gRNAs for targeting the *tbx16* locus and CFP in the lens for indication of the transgenes (Fig. 5A). Crosses between *Tg(piwil1:nCas9*^*ERT2*^*-n3U)* females and *Tg(U6:tbx16-gRNAs;cryaa:CFP)* males produced *Tg(piwil1:nCas9*^*ERT2*^*-n3U;U6:tbx16-gRNAs;cryaa:CFP)* triple transgenic embryos, which were treated with 40 μM 4-OHT from 4 hpf onwards (Fig. 5B). As observed at 30 hpf, the treated embryos looked morphologically normal, which were in sharp contrast to the spadetail-like *tbx16*^*tsu-G115*^ (designate allele code) mutants obtained by ENU-mediated mutagenesis in our lab (Fig. 5C). This result suggests that *tbx16* mutations might have rarely happened in somatic cells of 4-OHT treated transgenic embryos. Next, we picked up 7-8 PGCs from each 4-OHT treated 2 dpf embryo with mCherry expression in PGCs and genotyped each PGC after cloning and sequencing of the *tbx16* target sequence. Result revealed that 47.7% (21 of 44) of PGCs from 6 embryos carried *tbx16* mutant alleles, 23.8% (5 of 21) of which were biallelic mutations (Fig. 5D). Among the mutant alleles, the predominant one was the 31-bp deletion (65.2%) (Fig. 5E). These data indicate that stage-specific induction of nCas9^ERT2^-based gene editing activity also works for the *tbx16* locus in PGCs.

**Fig. 5.**
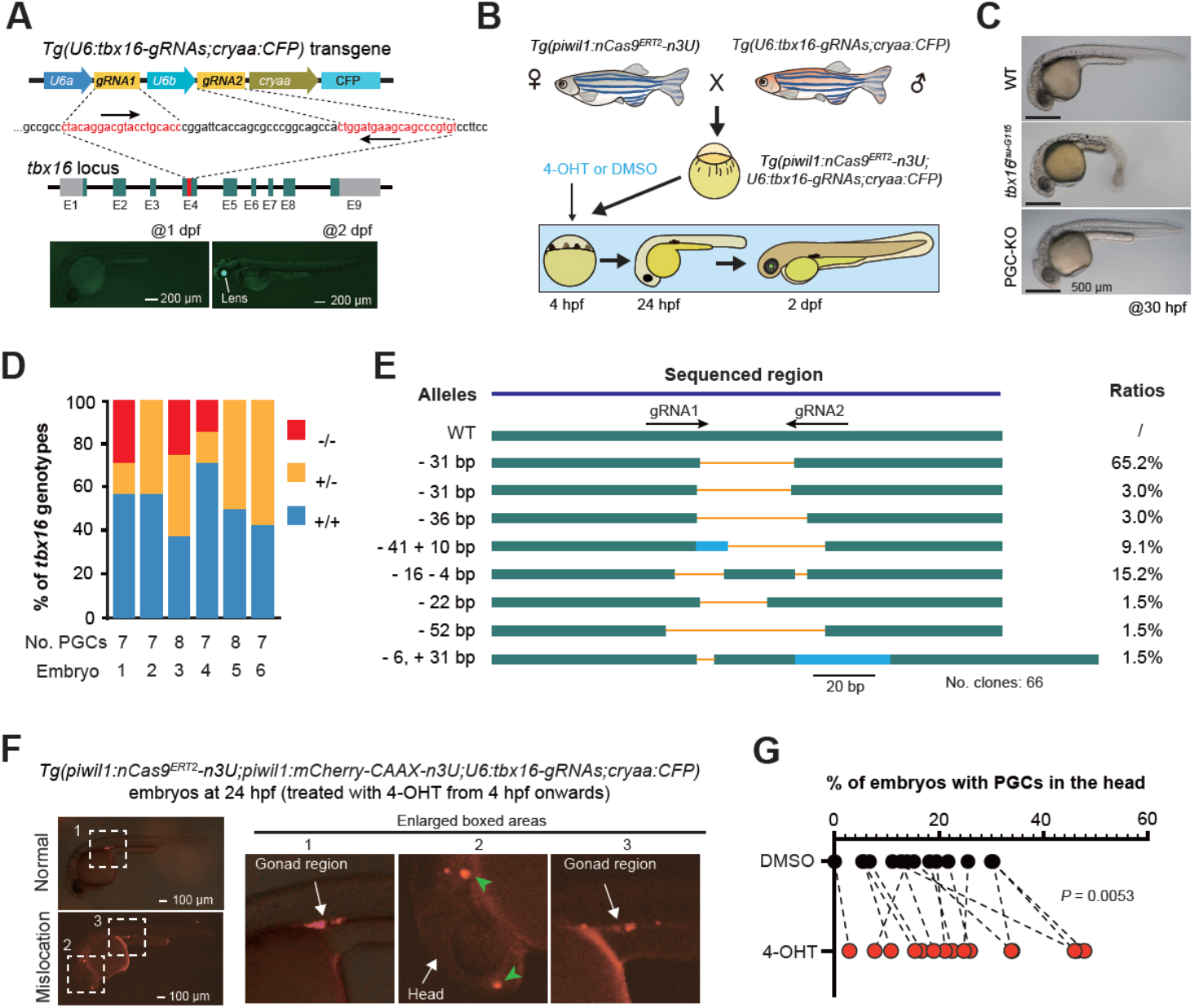
PGC-specific mutation of *tbx16* affects migration. (A) Genomic structure of the zebrafish *tbx16* locus and generation of *tbx16*-*gRNAs* expressing stable germline. The direction of *gRNA* recognition sequences was indicated by arrows. E1-E9, exons; gray areas, untranslated regions. The low panel showed the transgenic embryos with CFP expression in the lens at 2 dpf but not at 1 dpf. (B) Procedures to obtain embryos expressing nCas9^ERT2^ and *tbx16-gRNAs* and to treat the embryos. (C) Morphology of different types of embryos at 30 hpf. *tbx16*^*<sup>tsu-G115</sup>*^ mutant embryos carried biallelic mutations in all cells (whole-body knockout, Wb-KO) and exhibited spadetail-like phenotype while all *Tg(piwil1;nCas9*^*<sup>ERT2</sup>*^*-n3U;U6:tbx16-gRNAs;cryaa:CFP)* embryos treated with 4-OHT (PGC-KO) looked normal. (D) Ratio of PGC genotypes in PGC-KO embryos as in (B). Individual PGCs were isolated from individual 4-OHT treated *Tg(piwil1;nCas9*^*<sup>ERT2</sup>*^*-n3U;piwil1:mCherry-CAAX-n3U;U6:tbx16-gRNAs;cryaa:CFP)* embryos at 2 dpf and subjected to amplification, cloning and sequencing of the *tbx16* target region, allowing identification of each PGC’s genotype. −/−, two mutant alleles; +/−, one mutant allele; +/+, two WT alleles. (E) Allelic mutant types and ratios. Green, WT sequence; gray, untranslated regions; orange line, deleted region; blue, inserted region. (F) Examples of PGC locations in 24-hpf embryos. The boxed areas in the left panel were enlarged in the right panels. In the second boxed area, PGCs were apparently mislocated in the head region. (G) Ratio of embryos with PGCs in the head in 4-OHT treated embryos (as shown in (F) and DMSO-treated control embryos.

To look into the effect of *tbx16* mutations in PGCs, we observed PGC locations in *Tg(piwil1:nCas9*^*ERT2*^*-n3U;piwil1:mCherry-CAAX-n3U;U6:tbx16-gRNAs;cryaa:CFP)* embryos at 24 hpf by fluorescent microscopy, which were derived from crosses between *Tg(piwil1:nCas9*^*ERT2*^*-n3U;piwil1:mCherry-CAAX-n3U)* females and *Tg(U6:tbx16-gRNAs;cryaa:CFP)* males. The mCherry^+^-PGCs were normally well aligned in bilateral gonads at the anterior end of the yolk extension at 24 hpf; abnormal location of mCherry^+^-

PGCs occurred in the head or trunk region (Fig. 5F). The above transgenic embryos of each batch were divided into two groups, one of which was treated with 40 μM 4-OHT from 4 hpf onwards and the other one with DMSO. We then counted the number of embryos with obvious ectopic PGCs in the head region at 24 hpf. Results indicated that, on average, the 4-OHT group had a significantly higher proportion of embryos with ectopic PGCs (Fig. 5G). This result suggests that Tbx16 expressed in PGCs plays a role in pledging normal migration of PGCs during somitogenesis even though the involvement of Tbx16 in somatic cells in this process cannot be excluded.

### Inducible nCas9^ERT2^ system works in mouse early embryos

To explore the broad applicability of our nCas9^ERT2^ system in other species, we briefly tested blastomere-specific genome editing in mouse early embryos by targeting the *Cdx2* gene, which is specifically expressed in the trophectoderm (TE) of blastocysts and represses the expression of *Nanog* and *Oct4* in inner cell mass (ICM) (64). We co-injected synthetic *nCas9*^*ERT2*^ mRNA, *nls-mCherry* mRNA, and two *Cdx2* gRNAs in different combinations into one blastomere of late 2-cell stage embryos, immediately followed by treatment with 10 μM 4-OHT or DMSO (Fig. 6A). At 24 hours post treatment (at 8-cell stage), 4-OHT or DMSO were removed by wash, and embryos were either used for isolating mCherry-positive blastomeres for *Cdx2* amplification and sequencing or cultured in fresh medium to the blastocyst stage for cell lineage examination. Sequencing results of pooled mCherry-positive blastomeres of 8-cell stage embryos disclosed that mutant *Cdx2* alleles accounted for 58.82% (10/17) in blastomeres derived from embryos injected with all three RNA species and treated by 4-OHT (M group). In contrast, no mutant alleles were detected in blastomeres derived from embryos injected similarly but treated with DMSO (C3 group) (Fig. 6B). Genotyping individual mCherry-positive blastomeres revealed a predominance of biallelic mutations and different types of mutations (Fig. 6C and D) in M group embryos. Immunostaining of embryos at the blastocyst stage showed that the M group had a higher proportion of mCherry-positive blastomeres with loss of Cdx2 expression than the other control groups although the ratios of mCherry-positive cells per embryo were comparable among the M group and control groups (C1, C2, and C3) (Fig. 6E and F). Consistent with the idea that Cdx2 is a suppressor of *Nanog* transcription (64), strikingly, the percentage of Nanog^+^;mCherry^+^-blastomeres increased in the M group embryos (Fig. 6G and H). These observations together suggest that the inducible nCas9^ERT2^ system is applicable to mouse embryos for gene knockout.

**Fig. 6.**
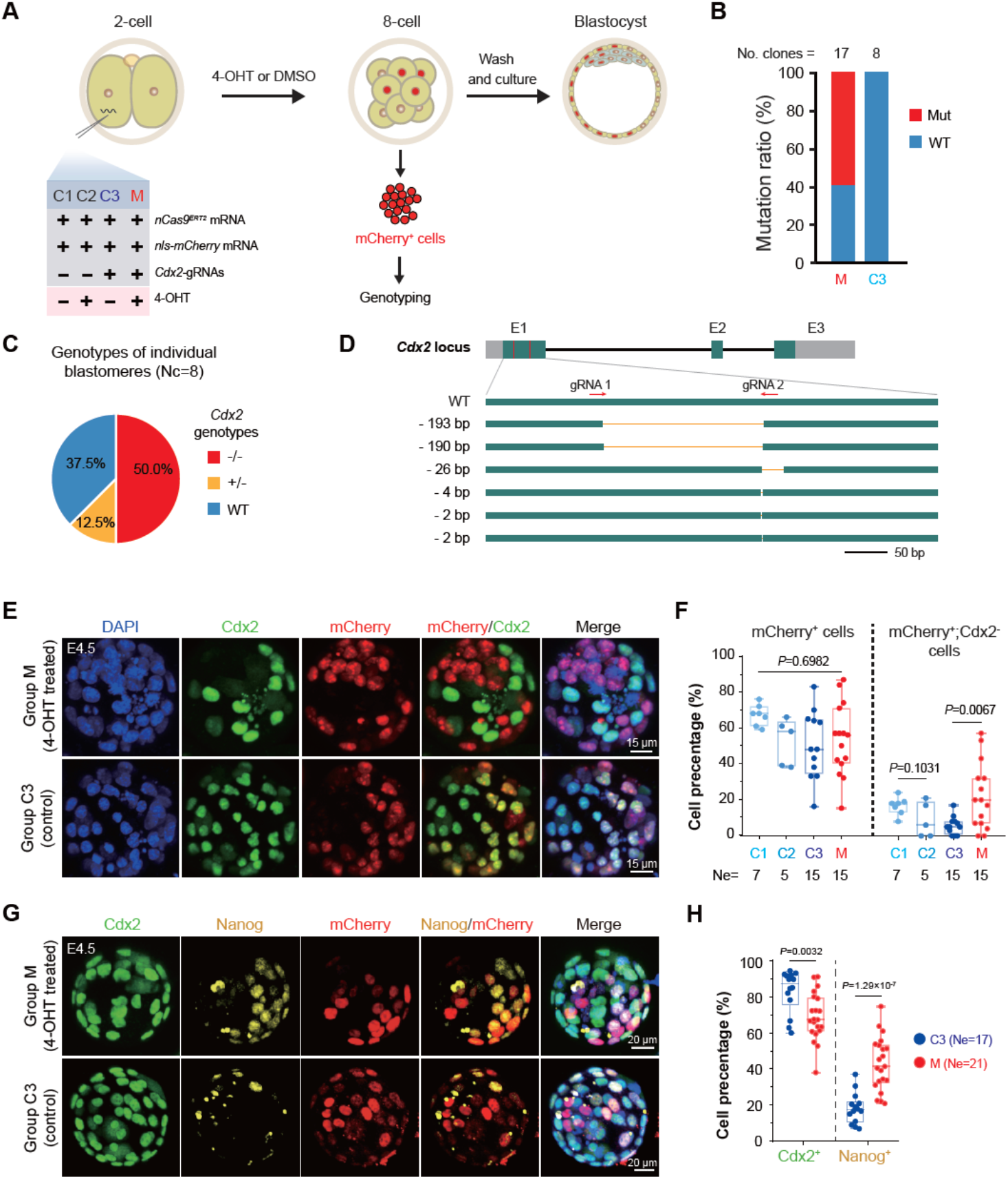
Mutation of *Cdx2* in selected blastomeres in mouse early embryos using inducible nCas9^ERT2^. (A) Experimental procedure. Different combinations (group C1-C3 and M) of indicated reagents were injected into one blastomere of 2-cell stage embryos. (B) Mutation ratios of *Cdx2* alleles in mCherry^+^ cells of Group M or Group C3 embryos at the 8-cell stage. A *Cdx2* region, including gRNA target sides, was amplified from pooled cells and cloned for sequencing. The number of sequenced clones was indicated. Mut, mutant alleles; WT, wildtype allele. (C) Ratios of cells with specific *Cdx2* genotypes in Group M embryos. The mCherry+ cells were isolated and genotyped individually. −/−, two mutant alleles; +/−, one mutant allele; WT, two WT alleles. Nc, the number of genotyped cells. (D) Structure of WT and mutant alleles as revealed in (C). E1-E3, exons; gray, untranslated regions; arrows, direction of *gRNA* recognition sites; orange line, deleted region. (E and F) Representative immunofluorescent images of Cdx2 and mCherry expression (E) and ratio of cells with or without indicated markers (F) in different groups of blastocysts at E4.5. (G, H) Representative immunofluorescent images of Cdx2, Nanog and mCherry expression (G) and ratio of cells with Cdx2 or Nanog expression (H) in Group M or C3 blastocysts at E4.5. In (F and H), the denominator was total number of cells in individual blastocysts; *P*-values of *t-test* (unpaired, two-sided) were indicated; Ne, the number of blastocysts.

## Discussion

In this study, we established an inducible gene editing approach to perform spatiotemporal gene knockout based on 4-OHT inducible nCas9^ERT2^. As a proof-of-concept, we applied this system to GC-specific gene knockout by expressing nCas9^ERT2^ in GCs driven by the GC-specific *piwil1* promoter, expressing gene-specific gRNAs ubiquitously, and stage-specific treating with 4-OHT.

Our inducible gene editing system is composed of two major components: a transgenic line with tissue-specific expression of 4-OHT inducible nCas9^ERT2^ as an acceptor and a transgenic line with ubiquitous expression of gene-specific gRNAs. The latter could be replaced by the injection of synthetic gRNAs. Once a tissue-specific transgenic line with strong nCas9^ERT2^ expression, it could be used for performing gene knockout of many genes, even for simultaneous knockout of several genes to study their functional redundancy. This is particularly useful in the zebrafish in which loxP knockin usually has a very low efficiency. In case several genes need to be knocked out simultaneously to study their functional redundancy, the application of the Cre/loxP system has to first generate at least one loxP knock-in line for each gene and then intercross them several times to obtain animals with multiple loxP knock-ins, which is usually time-consuming and often impractical. In contrast, the nCas9^ERT2^ system only needs one nCas9^ERT2^ transgenic line plus co-injection of several genes’ gRNAs for simultaneous knockout of these genes.

In our inducible system, nCas9^ERT2^ nuclear translocation relies on the action of 4-OHT. The concentration and treatment duration of 4-OHT should be optimized case by case. We noted that nCas9^ERT2^-mediated gene mutation does not occur in all cells of a target tissue, which may be ascribed to differential accessibility to the target gene locus in different cell types or cell statuses. Besides, only some target cells carry mutations in both alleles, which may make phenotypic analysis complicated. Nevertheless, the inducible nCas9^ERT2^ system would be very useful to explore stage-specific and/or tissue-specific function of a gene whose mutation in a whole organism may lead to death or pleiotropic effect.

## Materials and Methods

### Breeding of zebrafish

The WT Tübingen strain and all transgenic zebrafish lines used in this study were normally raised at 28.5°C in standard housing systems, with ethical approval from the Animal Care and Use Committee of Tsinghua University. Embryo developmental stages were defined as standard developmental stages (65). The generation of transgenic lines is described below.

### Construction of plasmids

In this work, we constructed three types of plasmids for mammalian cell transfection, transgenic zebrafish line construction, and *in vitro* transcription of mRNA, respectively. All of the transgenic plasmids in this research were constructed with *Tol2* transposon-based vectors. *SI Appendix Table S1* lists the plasmid information about vectors and restriction enzyme sites for insertions. *SI Appendix Table S2* presents the primers used for the inserted sequences in these plasmids, including four kinds of *piwil1* promoter, *cryaa* promoter, and *ef1a* promoter. The majority of the plasmids were constructed by the Gibson Assembly method (66), except for *pU6-EMX1 gRNA-Cas9-T2A-GFP*, which was ligated with T4 ligase following restriction enzyme cleavage. The *U6:gRNA* fragment in the *pU6-EMX1 gRNA* plasmid originated from *pU6-EMX1 gRNA-Cas9-T2A-GFP*.

### Cell culture and transfection

HEK293T cells were cultured in DMEM (Dulbecco’s modified eagle medium) supplemented with 10% FBS (fetal bovine serum) and PS (penicillin/streptomycin) at 37° C in a humidified incubator with 5% CO_2_. When the cell density is about 50%, plasmid transfection could be performed using VigoFect (T001, Vigorous). Plasmid transfection requires 2 μg per well in a 6-well cell culture plate. The mixture of transfection reagent and plasmids was obtained according to the manufacturer’s instructions and then added into the well. After 8 hours, replace with fresh DMEM.

### Generation of transgenic zebrafish lines

After plasmid construction, 30 pg of plasmid and 200 pg of *Tol2* transposase mRNA were co-injected into WT zygotes. For transgenic lines with fluorescence, embryos were screened out by observing fluorescence with fluorescent stereomicroscope (Olympus, MVX10) at 1 or 2 dpf. *Tg(piwil1:nCas9*^*ERT2*^*-n3U)* and *Tg(piwil1:nCas9-n3U)* transgenic embryos did not express fluorescence, so these embryos were screened by genotyping and *in situ* hybridization to detect *Cas9* mRNA expression.

### Mouse embryo culture

The mouse strain C57BL/6N was used in this study. Mice were maintained on a 12/12-hour light/dark cycle, 22-26°C, with sterile pellet food and water ad libitum. 12-14 g female mice were super-ovulated by intraperitoneal injection of 10 IU pregnant mare serum gonadotropin (PMSG, Sansheng Biological Technology, 21958956) and 10 IU human chorionic gonadotrophin (HCG, Sansheng Biological Technology, 110041282) at the interval of 47.5 hours. Then, the female mice were mated with 8 to 20-week-old male mice one to one. Embryos were collected from the oviduct of female mice with a vaginal plug in the morning, and granule cells were removed by digestion with 300 μg/mL hyaluronidase (Sigma, H4272) in M2 media (Merck Millipore, MR-015-D). Fertilized embryos with the second polar body were picked out and cultured in a 30 μL KSOM (Merck Millipore, MR-121-D) droplet covered by mineral oil (Sigma, M8410) in 37 °C incubator filled with 5% CO_2_. Developmental stages of mouse embryos were defined as follows: early 1-cell stage, 18-21 hpi (hours post HCG injection) or 9-12 hpf; late 2-cell stage, 48-50 hpi or 39-41 hpf.

### 4-OHT treatment of zebrafish and mouse embryos

The 4-OHT is dissolved in DMSO to 10 mM and stored at −20°C, kept out of the light. When treating zebrafish, 4-OHT is diluted with Holtfreter water (for embryos and larvae) or water from the housing system (for adults) to the proper concentration and kept away from light. When confirming whether the nCas9^ERT2^ protein could respond to 4-OHT treatment, embryos at the sphere stage were incubated in 6-well plates with 3 mL of 20 μM 4-OHT. When detecting the mutation efficiency of target genes, the embryos were incubated in 6-well plates with 3 mL of 40 μM 4-OHT to induce a higher mutation efficiency. The treatment began at the sphere stage and ended at 48 hpf. The young zebrafish at 45 dpf were incubated in 30 mL of 5 μM 4-OHT. When treating mouse embryos, 4-OHT is diluted with KSOM media to 10 μM and kept out of the light.

### Immunofluorescence staining of HEK293T cells, zebrafish embryos, and mouse embryos

HEK293T cells, zebrafish embryos, and mouse embryos were all fixed with 4% paraformaldehyde (Sigma, P6148) in PBS at 4 °C. The HEK293T cells were fixed for 30 minutes, whereas the zebrafish and mouse embryos were fixed overnight. Following fixation, zebrafish embryos were mechanically dechorionated and dehydrated into 100% methanol with a graded PBS: methanol series (3:1, 1:1, 1:3), and stored at −20°C, which are ready for immunofluorescence experiments. Stained HEK293T cells and embryos were imaged with an Olympus FV3000 microscope. The confocal data were processed using Imaris software.

Immunofluorescence staining was carried out as the standard protocol (68, 69). Briefly, HEK293T cells were permeabilized in 0.4% PBST (0.4% Triton X-100 in PBS) for 10 minutes, blocked with 3% BSA (3% bovine serum albumin in PBS) for 30 minutes, and treated with primary antibodies diluted in blocking buffer overnight at 4°C. The primary antibodies included mouse anti-Cas9 (Abcam, Ab191468, 1:200 dilution) and rabbit anti-mCherry (Easybio, BE2027, 1:200 dilution). The next day, HEK293T cells were incubated in DAPI (ThermoFisher, 1:10000 dilution) and fluorochrome-conjugated secondary antibodies in blocking buffer (1:200 dilution) for 2 hours, with all subsequent steps kept in the dark. The secondary antibodies included Alexa Fluor 488 AffiniPure Goat anti-Mouse IgG (H+L) (Jackson ImmunoResearch Labs, 115-545-003) and Rhodamine (TRITC) AffiniPure Goat Anti-Rabbit IgG (H+L) (Jackson ImmunoResearch Labs, 111-025-003).

Zebrafish embryos before 24 hpf were permeabilized in 0.5% PBST for 30 minutes, whereas embryos at 24 hpf were permeabilized in 1% PBST overnight. After that, embryos were treated for antigen retrieval with preheated 10 mM sodium citrate (pH 6.0), maintained at 95°C for 10 minutes, and cooled on a benchtop for at least 30 minutes. The embryos were then blocked and treated with primary and secondary antibodies diluted in blocking buffer, similar to the experiment methods for HEK293T cells, except that the incubation of DAPI and fluorochrome-conjugated secondary antibodies was overnight at 4°C. The primary antibodies included mouse anti-Cas9 (Abcam, Ab191468, 1:100 dilution), rabbit anti-Ddx4 (GeneTex, GTX128306-S, 1:1000 dilution), and rabbit anti-mCherry (Easybio, BE2027, 1:200 dilution). The secondary antibodies included Alexa Fluor 647 AffiniPure Goat anti-Rabbit IgG (H+L) (Jackson ImmunoResearch Labs, 111-605-003), Rhodamine (TRITC) AffiniPure Goat Anti-Rabbit IgG (H+L) (Jackson ImmunoResearch Labs, 111-025-003), and Alexa Fluor 488 AffiniPure Goat anti-Mouse IgG (H+L) (Jackson ImmunoResearch Labs, 115-545-003).

Mouse embryos were permeabilized in 0.5% PBST for 30 minutes, and the subsequent experiment procedures were similar to those of zebrafish embryos but without antigen retrieval. The primary antibodies included mouse anti-Cdx2 (BioGenex, MU392A-UC, 1:200 dilution), rabbit anti-Nanog (Abcam, Ab214549, 1:100 dilution), and rabbit anti-mCherry (Easybio, BE2027, 1:200 dilution). Using the same secondary antibodies as mentioned above.

### Design and test of gRNAs

The gRNAs of target genes in this study were designed on the CHOPCHOP website (https://chopchop.cbu.uib.no/), and the genotyping primers were also designed on this website. To test the gRNA mutation efficiency, 300 pg of gRNA and 200 pg of *Cas9* mRNA were co-injected into WT zygotes. *Cas9* mRNA was zebrafish codon-optimized and transcribed *in vitro* with the mMESSAGE mMACHINE T7 Transcription kit (ThermoFisher, AM1344), and gRNAs were transcribed *in vitro* with the MEGAscript T7 Kit (ThermoFisher, AM1333). The mutation efficiency of gRNA was determined by genotyping and T7E1 assay with injected embryos at 24 hpf. *SI Appendix Table S2* lists the primers used for gRNA generation.

### Collection of zebrafish PGCs and somatic cells, and mouse blastomere

For 24 hpf and 2 dpf zebrafish embryos, PGC-containing regions of embryos were excised and transferred into 1.5 mL Eppendorf tubes. 10 to 50 μL of trypsin was added per tube, and tubes were placed at 37°C for digestion. The mixture was pipetted 20 times through 200 μL tips every 10 minutes until no visible tissue remained. Cell suspensions were then transferred to clear and RNase-free Petri dishes and mCherry-labeled PGCs were manually picked out under a fluorescence microscope with a mouth pipette. mCherry-negative cells are somatic cells utilized to determine if the target genes are mutated in somatic cells.

For 8-cell stage mouse embryos, the ZP (zona pellucida) of embryos was eliminated using acid Tyrode’s solution (Merck Millipore, MR-004-D). Then, the embryos were immediately transferred to 0.25% Trypsin (AMRESCO, 0458) at 37°C, separated into single blastomeres by gentle pipetting, and washed three times in KSOM media, which took approximately 5 to 10 minutes. Blastomeres were transferred to clear KSOM media, and mCherry-positive blastomeres were manually picked out with a mouth pipette under an Olympus inverted fluorescence microscope.

### Genome extraction and genotyping

For genome extraction of HEK293T cells, about 20,000 cells were collected and lysed in 15 μL of 50 mM NaOH at 95°C for 10 minutes, followed by the addition of 1/10 volume of 1 M Tris-HCl (pH 8.0) and vertexing. The same procedure was employed to extract the genomes of zebrafish and mouse embryos. 24 hpf zebrafish embryo was lysed with 30 μL of 50 mM NaOH, while the mouse blastocyst was lysed with 10 μL of 50 mM NaOH. The genome of adult zebrafish was extracted using the same method with clipped caudal fin tissue. The lysate would then be used as a genome template for PCR amplification. The genomes of PGC mixtures, individual PGCs, and individual mouse blastomeres were extracted and amplified with the Single Cell WGA Kit (Vazyme, N603), diluted with ddH_2_O to a suitable concentration, and used as PCR templates. *SI Appendix Table S2* summarizes the primer sequences for genotyping.

When detecting the target gene mutations in HEK293T cells, embryos, and cell mixtures, PCR products were further examined by T7EI (T7 Endonuclease I) (NEB) digestion at 37°C for 30 minutes, followed by sequencing for determination of the mutated sequence. The mutant gene would result in double bands after T7EI digestion and dual peaks in the Sanger sequencing map. To calculate the allelic mutation ratios, the PCR products of gRNA-targeted regions were ligated to the pBM18A vectors (Biomed, CL073) for 2 hours at room temperature, then transformed into competent *E*.*coli*. The next day, monoclones were picked out for Sanger sequencing. When genotyping a single PGC or mouse blastomere, the PCR products were also ligated to the pBM18A vectors for monoclonal Sanger sequencing to determine whether the PGC was hetero-mutated or homo-mutated.

### Statistical analysis

An average from multiple samples was expressed as mean ± SD (standard deviation). The significance of the difference between groups was analyzed by Student’s *t*-test (two-tailed). Significant levels were indicated in the corresponding context.

## Acknowledgments

We thank all members of the Meng lab for their intellectual and technical support. We would like to express our gratitude to Dr. Ming Shao for providing *pU6:gRNA* plasmids. We are grateful to the Cell Biology Facility and Sharing Core Facility affiliated with the Center of Biomedical Analysis, Tsinghua University, for technical assistance and daily equipment support.

## Funding

This work is financially supported by the National Natural Science Foundation of China (#31988101 to A.M.), the National Key Research and Development Program of China (#2023YFA1800300 to X.W. and 2018YFC1003304 to A.M.), and the Yunnan

Provincial Science and Technology Project at Southwest United Graduate School (#202302AO370011 to A.M.).

## Author contributions

Y. Li performed most experiments. W. Zhang and Z. Wei helped to generate and raise *Tg(piwil1:nCas9*^*ERT2*^*-n3U)* and *Tg(piwil1:nCas9-n3U)* transgenic lines. H. Li helped with mouse embryo immunostaining. X. Liu helped with experiments in HEK293T cells. T. Zheng and T. Aziz participated in some experiments and discussion. C. Xing participated in the model design. Y. Li, X. Wu, and A. Meng designed the study and prepared the manuscript.

## Competing interests

The authors declare that they have no competing interests.

---

**Fig. S1.**
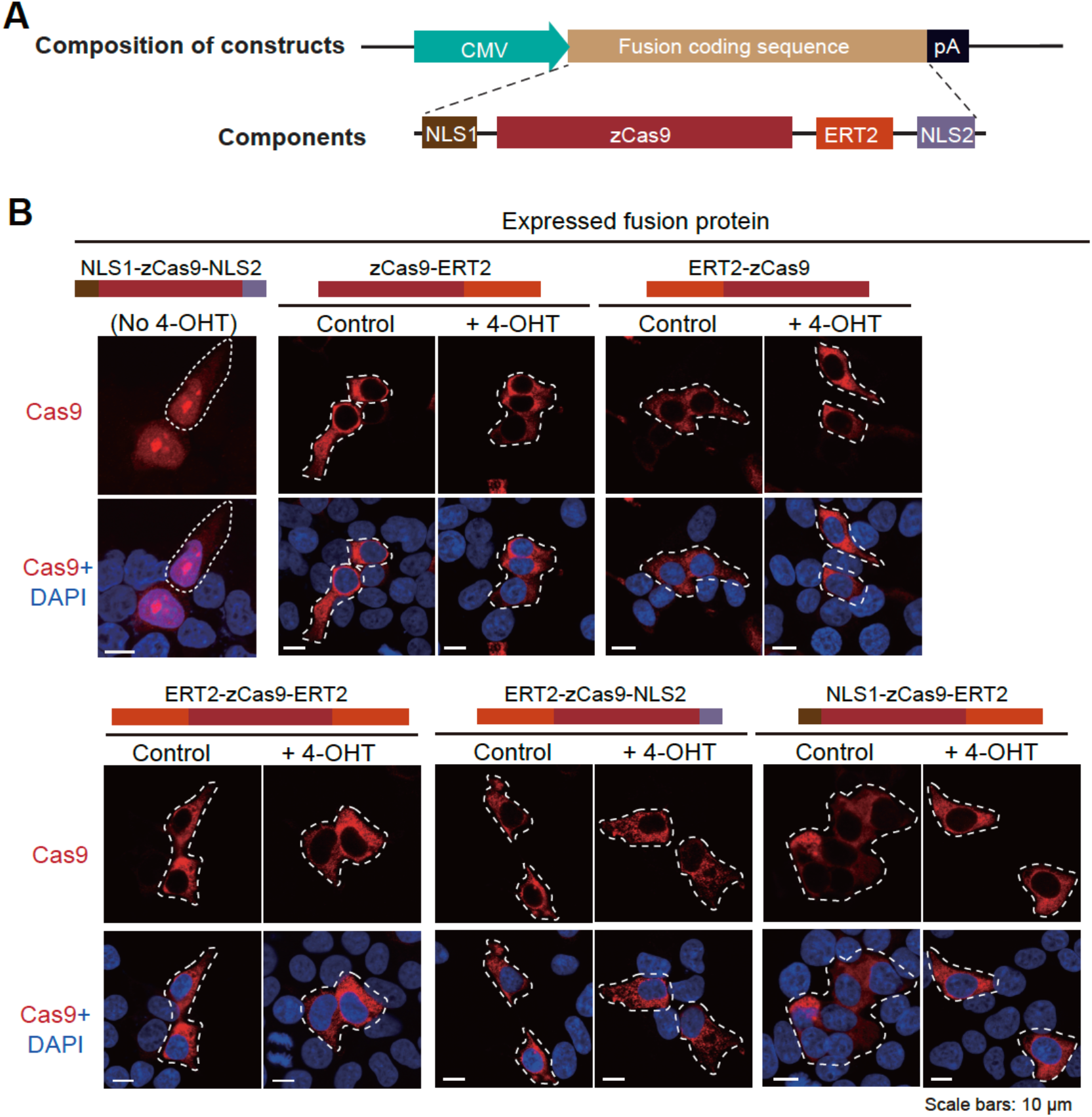
Test of nuclear translocation ability of Cas9 protein expressed in HEK293T cells. (A) General structure of transfection constructs. CMV, Cytomegaloviral promoter; zCas9, Cas9 with codon optimization for the zebrafish; ERT2, human estrogen receptor with three missense mutations (G400V/M543A/L544A) in the ligand binding domain; NLS1, SV40 nuclear localization signal, NLS2, nucleoplasmin NLS; pA, polyadenylation signal. (B) Nuclear translocation ability of different Cas9 fusion proteins with or without 4-OHT treatment. The transfected cells were immunostained with Cas9 antibody plus DAPI staining for nuclei 24 h posttreatment. The top of each group of images showed the schematic structure of the Cas9 fusion protein.

**Fig. S2.**
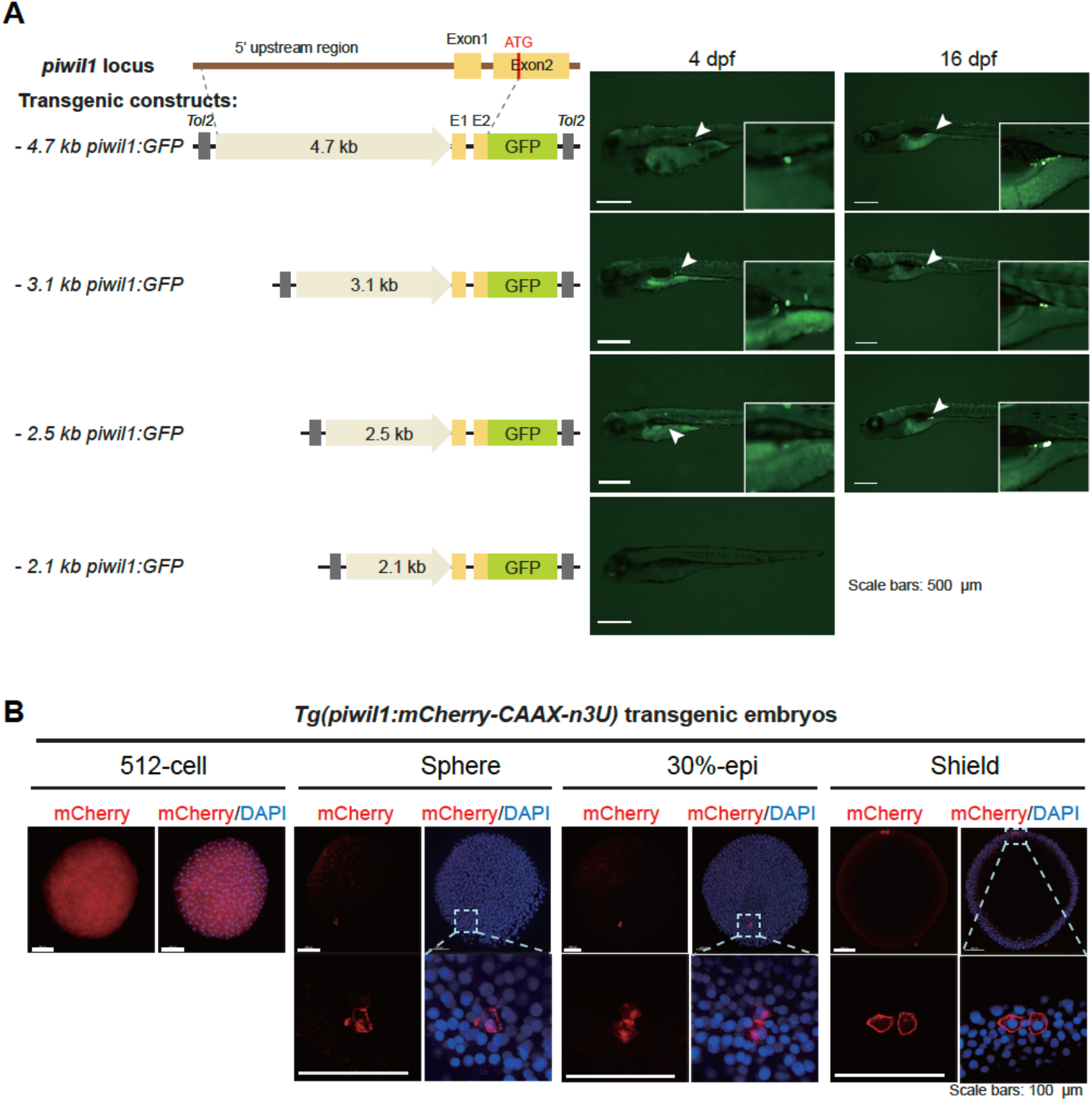
Test of GC specificity of the zebrafish *piwil1* promoter. (A) Identification of shorter *piwil1* promoter with GC specificity. Left, *Tol2* transposon-based construct structures; right, representative GFP expression patterns in zebrafish larvae after injection with corresponding constructs. GFP was observed under fluorescent dissection microscopy. Note that the 2.5-kb *piwil1* promoter retained GC-specific transcription activity and was used subsequently. (B) A transgenic line with GC-specific expression of membrane-localizing mCherry driven by the 2.5-kb *piwil1* promoter. CAAX, cell membrane localizing signal; n3U, 3’ untranslated region of the zebrafish *nanos3* gene. Note that mCherry was ubiquitously expressed at the 512-cell stage but came restricted to PGCs thereafter. The boxed areas in the top panel were enlarged in the low panel.

**Table S1.** Information of plasmids.

| Plasmid | Backbone | Restriction enzyme sites for insertions |  |  |
| --- | --- | --- | --- | --- |
| *pCS2-zCas9-ERT2* | pCS2 | BamHI / EcoRI |  |  |
| *pCS2-ERT2-zCas9* |  |  |  |  |
| *pCS2-ERT2-zCas9-ERT2* |  |  |  |  |
| *pCS2-ERT2-zCas9-NLS2* |  |  |  |  |
| *pCS2-NLS1-zCas9-ERT2* |  |  |  |  |
| *pCS2-nCas9<sup>ERT2</sup>* |  |  |  |  |
| *pCS2-nCas9* |  |  |  |  |
| *pU6a-EMX1-gRNA* |  |  |  |  |
| *pU6-EMX1-gRNA;*<br>*CMV-NLS1-Cas9-NLS2-T2A-GFP* | PX458 | BbsI |  |  |
| *-4.7 kb piwill:GFP* | pSTG | EcoRI / BamHI |  |  |
| *-3.1 kb piwill:GFP* |  |  |  |  |
| *-2.5 kb piwill:GFP* |  |  |  |  |
| *-2.1 kb piwill:GFP* |  |  |  |  |
| *piwill:mCherry-CAAX-n3U* | pSTG | EcoRI / NotI |  |  |
| *piwill:nCas9<sup>ERT2</sup>-n3U* |  |  |  |  |
| *piwill:nCas9-n3U* |  |  |  |  |
| *U6:tbx16-gRNAs;cryaa:CFP* |  |  |  |  |
| *U6:hwa-gRNAs;efla:GFP* |  |  |  |  |
| Plasmid | Vector | Restriction enzyme sites for insertions | Restriction enzyme sites for *in vitro* transcription | Transcriptase |
| *pXT7-nCas9* | pXT7 | EcoRI / SpeI | BamHI | T7 |
| *pXT7-nCas9<sup>ERT2</sup>* |  |  |  |  |

**Table S2.**
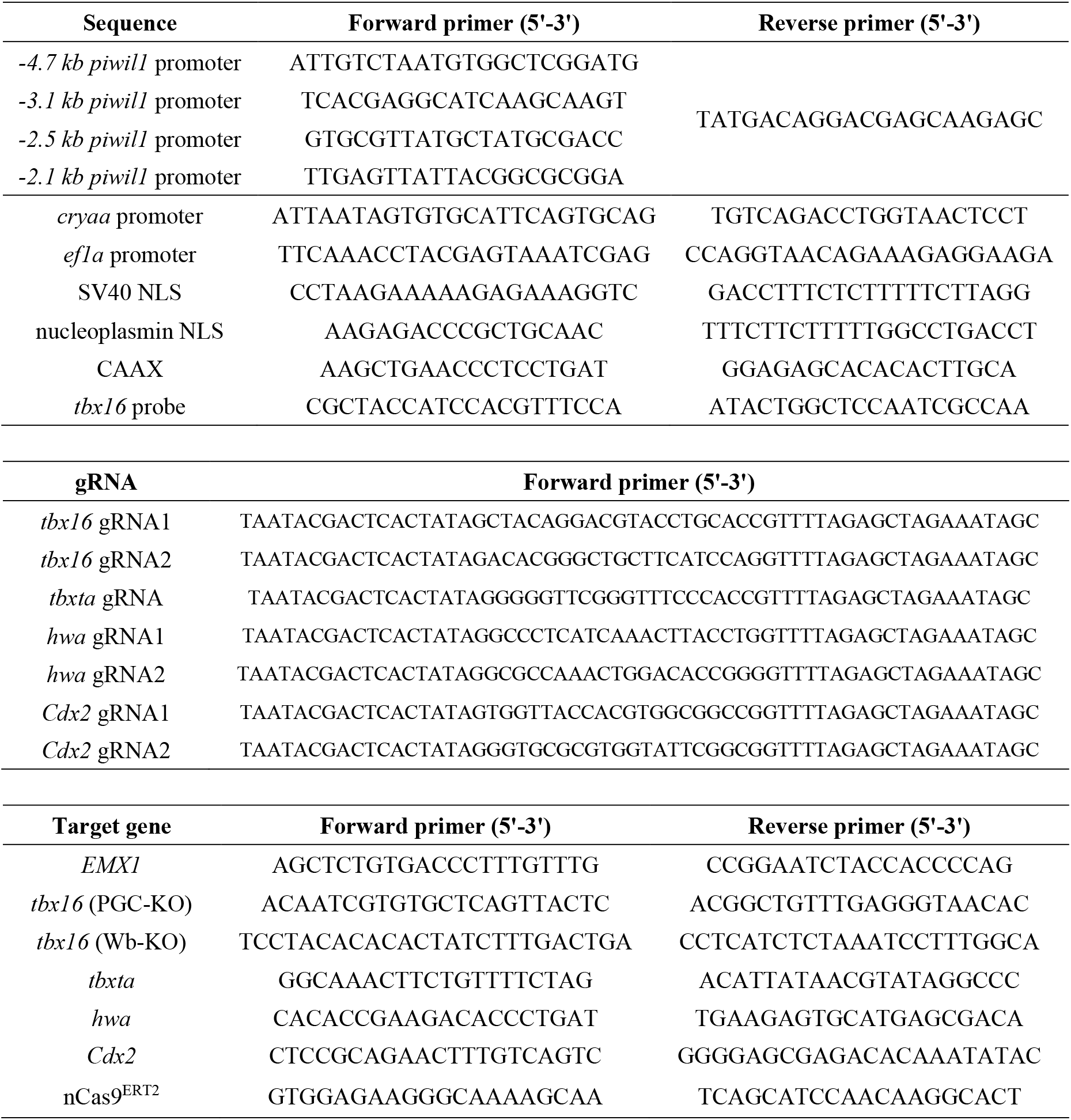
Information of primers.

## Notes

### Competing Interest Statement

The authors have declared no competing interest.

